# Novel monoclonal antibodies showing broad neutralizing activity for SARS-CoV-2 variants including Omicrons BA.5 and BA.2.75

**DOI:** 10.1101/2022.09.02.506305

**Authors:** Hanako Ishimaru, Mitsuhiro Nishimura, Lidya Handayani Tjan, Silvia Sutandhio, Maria Istiqomah Marini, Gema Barlian Effendi, Hideki Shigematsu, Koji Kato, Natsumi Hasegawa, Kaito Aoki, Yukiya Kurahashi, Koichi Furukawa, Mai Shinohara, Tomoka Nakamura, Jun Arii, Tatsuya Nagano, Sachiko Nakamura, Shigeru Sano, Sachiyo Iwata, Yasuko Mori

## Abstract

We identified novel neutralizing monoclonal antibodies against SARS-CoV-2 variants (including Omicron) from individuals received two doses of mRNA vaccination after they had been infected with wildtype. We named them MO1, MO2 and MO3. MO1 shows high neutralizing activity against authentic variants: D614G, Delta, BA.1, BA.1.1, BA.2, and BA.2.75 and BA.5. Our findings confirm that the wildtype-derived vaccination can induce neutralizing antibodies that recognize the epitopes conserved among the SARS-CoV-2 variants (including BA.5 and BA.2.75). The monoclonal antibodies obtained herein could serve as novel prophylaxis and therapeutics against not only current SARS-CoV-2 viruses but also future variants that may arise.

## Introduction

Severe acute respiratory syndrome-coronavirus 2 (SARS-CoV-2) caused the explosive Coronavirus disease 2019 (COVID-19) pandemic that began in 2019; SARS-CoV-2 has infected over 577 million people worldwide and is responsible for more than 6.4 million deaths as of August 4, 2022 (https://covid19.who.int/). The Omicron variant (BA.1; B.1.1.529) of SARS-CoV-2 was first reported in South Africa in November 2021 and had spread worldwide (https://www.cdc.gov/coronavirus/2019-ncov/variants/Omicron-variant.html) (Dejnirattisai et al., 2022). The spread of the Omicron variant has caused an increase of COVID-19 infections, and the Omicron BA.5 variant has spread worldwide (Tegally et al., 2022). We and other research groups have shown that three doses of the mRNA vaccine can protect individuals against the BA.1 and BA.2 variants of SARS-CoV-2 (Muik et al., 2022) (Furukawa et al., 2022) (Tjan et al., 2022). We have also observed that neutralizing antibodies against Omicron variants are increased in the sera of individuals received a two-dose mRNA vaccine after they had been infected with wildtype SARS-CoV-2 (D614G) (Kurahashi et al., 2022a). We therefore screened and searched those individuals’ B cells, as we suspected that it is possible that these B cells could produce antibodies that have broad neutralizing activity against SARS-CoV-2 variants including Omicron BA.5. In the present study, we identified three novel monoclonal antibodies (mAbs) that are broadly effective against SARS-CoV-2 variants including Omicron BA.5.

## RESULTS

### B cells that produce antibodies with broad neutralizing activity against SARS-CoV-2 were obtained from the PBMCs of SARS-CoV-2-infected, and subsequently two-dose-vaccinated patients

To isolate mAbs targeting common epitopes of SARS-CoV-2 variants, we searched for antibody genes from PBMCs of infected and subsequently vaccinated patients. We selected three PBMCs from the patients who showed high neutralizing activity against all of the D614G, Delta, and Omicron BA.1 variants previously we reported (Kurahashi et al., 2022a). In the single B cells isolated from the PBMCs, ten antibody genes were further selected based on the ELISA findings of Fabs produced by the Ecobody technology (Ojima-Kato et al., 2017), and we tested the purified recombinant IgG antibodies with the V genes. Ten mAbs were examined for neutralizing activity, and we used the three (which we named MO1, MO2, and MO3) that showed neutralizing activity in the present study.

### Broadly neutralizing mAbs against SARS-CoV-2 variants

As depicted in Figure 1, we observed that the three neutralizing mAbs for D614G, i.e., MO1, MO2, and MO3, they recognize the SARS-CoV-2 spike protein of D614G. MO1 recognized the spike protein of five variants: D614G, Delta, BA.1, BA.2 and BA.5. MO2 recognized the spike protein of four of these variants (not BA.5). MO3 recognized three of the variants (not Delta or BA.5). We next assessed whether these three mAbs had neutralizing activity against SARS-CoV-2 variants. As shown in Figures 1B, C and D, the mAb MO1 inhibited all five variants and BA.1.1 variant with the following IC_50_ values: D614G (23.62 ng/mL), Delta (15.84 ng/mL), BA.1 (4.0 ng/mL), BA.1.1 (10.64 ng/mL), BA.2 (20.31 ng/mL), and BA.5 (15.67 ng/mL). MO2 inhibited four variants (BA.5 was the exception), with the following IC_50_ values: D614G (65.81 ng/mL), Delta (88.24 ng/mL), BA.1 (17.71 ng/mL), BA.1.1 (36.05 ng/mL) and BA.2 (151.2 ng/mL). MO3 suppressed three variants (Delta and BA.5 were the exceptions), with the following IC_50_ values: D614G (231.57 ng/mL), BA.1 (594.63 ng/mL), and BA.2 (701.95 ng/mL). Furthermore, all of three mAbs could neutralize BA.2.75 as well (data not shown).

**Figure 1.**
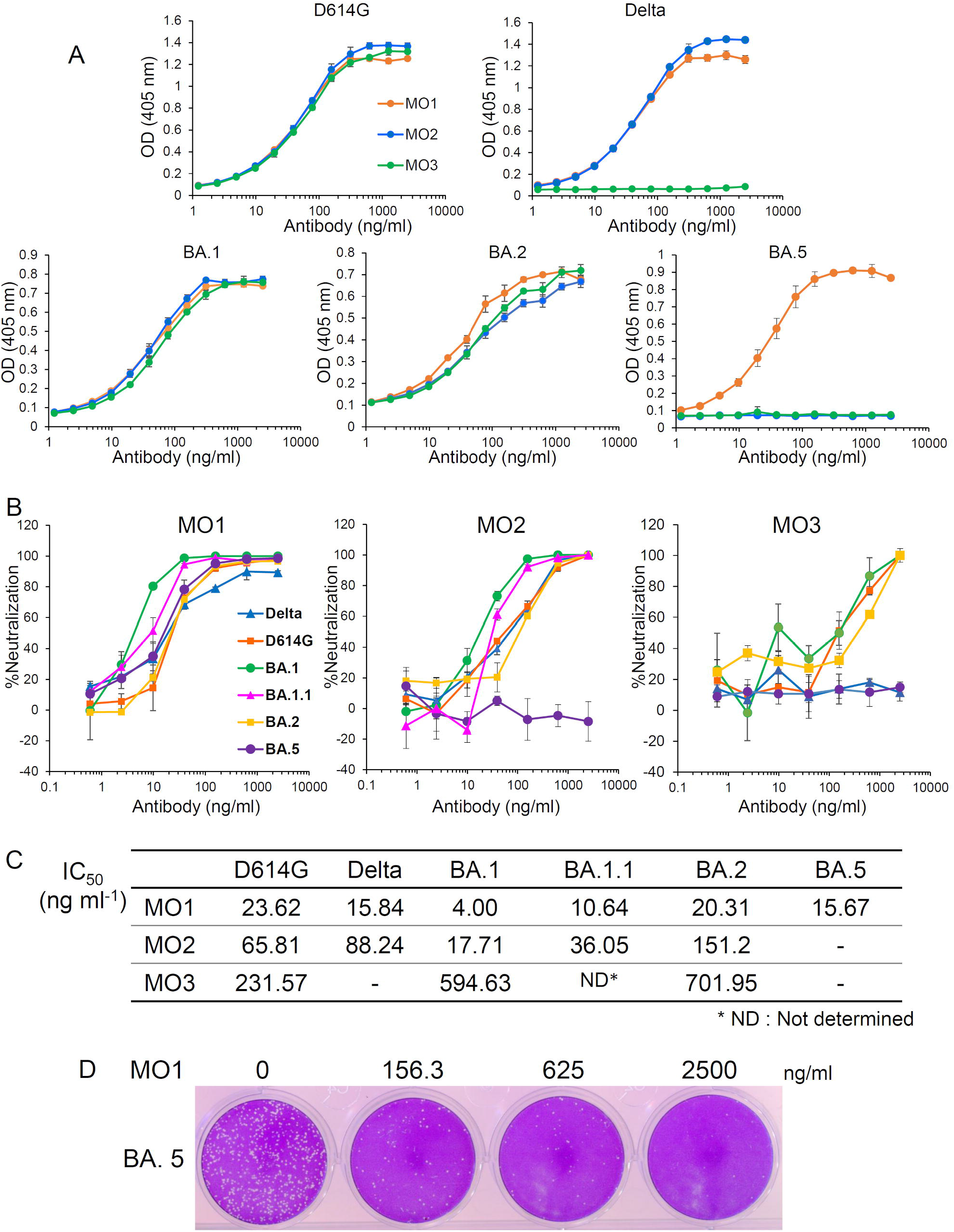
Identification of broadly neutralizing mAbs against SARS-CoV-2 variants. **(A)** The binding of three mAbs to the SARS-CoV-2 spike ectodomains of the D614G, Delta, BA.1, BA.2, and BA.5 variants as revealed by ELISAs. **(B)** The neutralizing activity of the three mAbs, MO1, MO2 and MO3 against D614G, Delta, BA.1, BA.1.1, BA.2 or BA.5 as evaluated by the plaque reduction neutralization test (PRNT). **(C)** The 50% inhibitory concentrations (IC50) of the mAbs MO1, MO2 and MO3 against the SARS-CoV-2 variants calculated from the above neutralization data (B) are shown. **(D)** A presentation of plaque reduction in MO1’s PRNT test against BA.5.

### The novel neutralizing mAb MO1 has high affinity against the SARS-CoV-2 BA.5 spike protein

The affinity between the SARS-CoV-2 BA.2 spike protein and the MO1 and MO2 mAbs was next evaluated by the Bio-layer interferometry (BLI) method. The prefusion-stabilized BA.2 spike ectodomain trimer hardly dissociated from MO1 or MO2, indicating high avidity due to multivalent binding sites on the spike trimer (data not shown). Both MO1 and MO2 showed high affinity with the BA.2 spike RBD, with the dissociation constants (K_D_) of 3.3 nM and 2.0 nM, respectively (Figures 2A, C). MO1 also has high affinity with the BA.5 spike RBD with the K_D_ of 11 nM (Figures 2B, C), whereas MO2 showed no binding to the BA.5 spike RBD, which is consistent with its loss of the neutralizing activity for BA.5.

**Figure 2.**
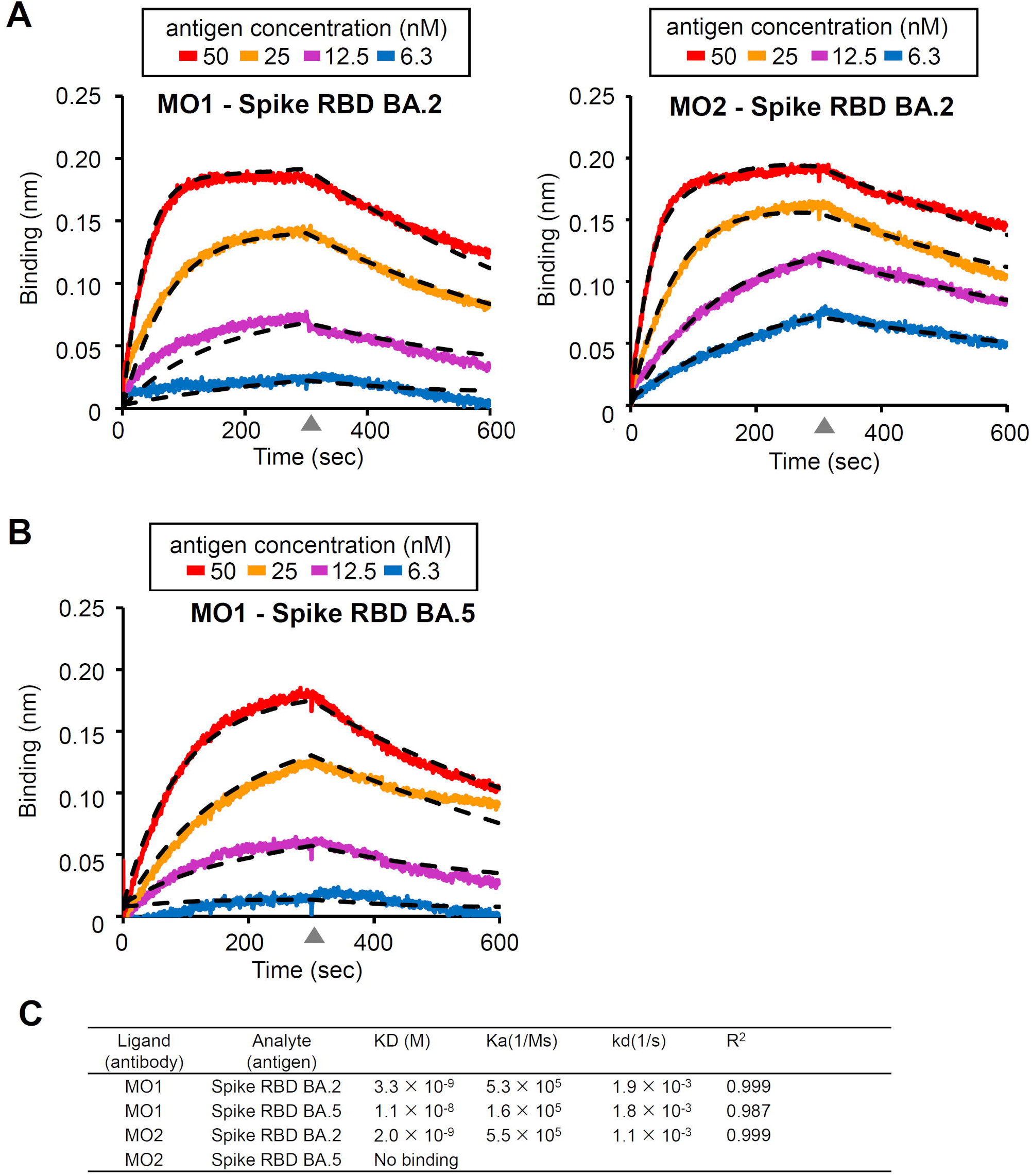
Analysis of the affinity between the three novel mAbs and spike antigens by biolayer interferometry (BLI). **(A)** The sensorgram for the BA.2 spike RBD’s binding to the mAb MO1 or MO2. *Dashed lines:* the fitting curves. **(B)** The same BLI analysis as in (A) between the BA.5 spike RBD and MO1. **(C)** Summary of the BLI kinetics evaluated from the curve fitting.

## DISCUSSION

The emergence of SARS-CoV-2 variants has prolonged the COVID-19 pandemic. The novel neutralizing mAb that we identified in the present study and named MO1 has strong neutralizing potency against all of the major SARS-CoV-2 variants described to date, including the Omicron BA.5 variant that is raging worldwide. The Omicron variants BA.1 and BA.2 are able to escape from the majority of the known neutralizing antibodies (Ai et al., 2022; Yamasoba et al., 2022a), and the escape ability of BA.5 is strengthened due to additional mutations such as F486V around the binding site of class 1 antibodies and the mutation site L452R around the binding site of class 3 antibodies as reported (Cao et al., 2022). Indeed, the neutralizing antibody MO2 identified herein, which can strongly neutralize both BA.1 and BA.2, lost neutralizing activity for BA.5. In contrast, MO1 has maintained neutralizing activity against BA.5. A conserved epitope avoiding the position of the L452R or F486V mutation sites is expected for MO1, considering the broad neutralizing activity of MO1 (Figure 1). Structural analysis by cryoelectron microscopy (cryo-EM) is under progress and the details of the MO1 epitope are under investigation.

It has been reported that the antibodies bebtelovimab (LY-CoV1404 (Westendorf et al., 2022)) and cilgavimab (AZD1061 (Dong et al., 2021)) have neutralizing activity against Omicron BA.1, BA.2, and BA.5, and that cilgavimab showed reduced activity against BA.5 (Yamasoba et al., 2022b). The binding affinity and neutralizing function of MO1 to BA.5 are high (Figures 1, 2). Our preliminary cryo-EM analysis indicated that MO1 accesses to the RBD from the similar side as the bebtelovimab and cilgavimab, while the binding site is substantially different from them.

MO1 is derived from PBMCs of individuals who were infected with a SARS-CoV-2 variant□—□presumably the wildtype with only the D614G mutation in the spike□—□and then received two doses of an mRNA vaccine encoding the spike gene of the ancestral SARS-CoV-2 (Kurahashi et al., 2022a). Our finding that MO1 can broadly neutralize the early SARS-CoV-2 variants and the Omicron BA.1, BA.1.1, BA.2 and BA.5 variants demonstrates that immunity against the spike with the ancestral SARS-CoV-2 RBD sequence can protect humans by inducing neutralizing antibodies, like MO1, that recognize conserved epitopes. Indeed, our serological study regarding the sustainability of the neutralizing antibodies after early SARS-CoV-2 (D614G) infection indicated that cross-reactive neutralizing antibodies are sustained for >6 months, although neutralizing antibodies specific for SARS-CoV-2 (D614G) decline (Kurahashi et al., 2022a; Kurahashi et al., 2022b). It is also noteworthy that the booster dose (three-dose) vaccination using the mRNA vaccine based on wildtype SARS-CoV-2 can induce neutralizing antibodies against Omicron BA.1 and BA.2 (Furukawa et al., 2022; Tjan et al., 2022).

It is speculated that MO1 recognizes a conserved epitope shared among SARS-CoV-2 variants considering its broad neutralizing activity. Isolation of MO1 from human memory B-cells may demonstrate that target sites remain available for neutralizing antibodies even in the spike of Omicron variants. Thus, the key task to develop effective immunity against a broad range of SARS-CoV-2 variants is thus to determine how to effectively induce antibodies against the conserved epitope. One of our previous studies revealed that patients who were infected with wildtype SARS-CoV-2 and then received a two-dose mRNA vaccination acquired high neutralizing antibody titers against Omicron BA.1 (Kurahashi et al., 2022a), and the neutralizing antibodies that were isolated in the present study were derived from donors with this background. Repeated exposure to the SARS-CoV-2 spike protein, irrespective of the variation in the sequence, may induce broadly neutralizing antibodies.

Cao et al. reported that BA.1-derived vaccine boosters may not achieve broad-spectrum protection against new Omicron variants because Omicron may evolve mutations to evade the humoral immunity elicited by BA.1 (Cao et al., 2022). In addition, all antibodies were found to be able to neutralize BA.2.75 variant newly appeared (Tan et al., 2022). The MO1 binding site observed by preliminary cryoelectron micrography is distant from mutation sites of the BA.2.75 variants (data not shown).

Our present findings demonstrate that the booster vaccination for the wildtype can induce neutralizing antibodies that may recognize common epitopes that are conserved among SARS-CoV-2 variants. It can also be speculated that a three- or four-dose vaccination based on the wildtype may stimulate memory B cells that can produce antibodies that have common epitopes conserved, showing broad neutralizing activity, and that booster vaccinations should thus be required even for previously infected individuals. The three new monoclonal antibodies identified in the present study could serve as novel therapeutics against not only the current SARS-CoV-2 variants but also new variants that arise in the future.

## Acknowledgments

We thank Kazuro Sugimura MD, PhD (Superintendent, Hyogo Prefectural Hospital Agency and Professor, Kobe University) for his full support of this study. We express our gratitude for the cooperation of researchers of the Division of Respiratory Medicine, Department of Internal Medicine, Kobe University Graduate School of Medicine, and medical staff of Hyogo Prefectural Kakogawa Medical Center. We thank BIKEN Innovative Vaccine Research Alliance Laboratories for providing the SARS-CoV-2 B2 strain used as the D614G variant herein. We also thank the National Institute of Infectious Disease Japan for providing the SARS-CoV-2 of Pango lineages AY.122, BA.1.18, BA.1.1, BA.2, BA.2.75 and BA.5 used here as the Delta, Omicron (B.1.159) BA.1, BA.1.1, BA.2, BA2.75 and BA.5 variants, respectively. This work is partially supported by Research Support Project for Life Science and Drug Discovery (Basis for Supporting Innovative Drug Discovery and Life Science Research: BINDS) from AMED under Grant Number JP22ama121001. YK was supported by the Taniguchi Memorial Scholarship program provided by the BIKEN Foundation. SS, LHT, MI, and GB were supported by Japanese Government (Monbukagakusho: MEXT) Scholarships.

## Funding

This work was supported by the Hyogo Prefectural Government and a grant from the Kobayashi Foundation.

## Author contributions

Conceptualization, HI, MN, LHT and YM.; Methodology, HI, MN, HS, KK and YM.; Sample collection: TN, SN, SS and SI.; Formal analysis, HI, MN, LHT, SS, MIM, GBE, HS, KK, NH, KA, YK, KF, MS TN, JA and YM.; Funding acquisition: YM.; Project administration, HI, MN, YM.; Supervision, YM.; Writing – original draft.; HI, MN and YM.; Writing – review & editing, HI, MN and YM.

## Declaration of interests

The authors declare no conflicts of interest in this research.

## Materials and Methods

### Collection of human samples

Blood samples were collected at Hyogo Prefectural Kakogawa Medical Center (Kakogawa, Japan) from patients who were infected with COVID-19 during the 5-month period of July–November 2020 and then received a two-dose messenger (m)RNA vaccine. In our earlier investigation (Kurahashi et al., 2022a), we identified the sera that had high neutralizing activity against SARS-CoV-2, and in the present study we used those patients’ (donors’) sera and data.

### Isolation of human antibodies’ genes from the donors’ PBMCs

In order to isolate human antibody genes from the above-described donor PBMCs, the sera’s memory B cells that expressed anti-SARS-CoV-2 spike Abs were screened and then sorted as single cells. Next, the anti-spike reactivity of the antibody Fab domain expressed from each V gene as shown by Ecobody technology (Ojima-Kato et al., 2017) was evaluated by an ELISA (enzyme-linked immunosorbent assay) (iBody, Nagoya, Japan).

The sequences of the selected V genes were determined and then subcloned into human immunoglobulin (IgG) mAb-expressing vectors. We used a capillary electrophoresis (CE) sequencer (model DS3000, Hitachi High-Tech, Tokyo) to confirm the sequences.

### Antibody expression and purification

With polyethyleneimine and the two plasmids containing the heavy-chain and light-chain sequences of the antibodies, recombinant mAbs were expressed in HEK293T (human kidney) cells by transfection.

For the purification of the antibodies, we added rProtein A Sepharose^®^ (Cytiva, Marlborough, MA) to the culture supernatant. The trapped mAbs was eluted with sodium citrate buffer (i.e., 40 mM trisodium citrate, pH 3.4). The solution was then immediately neutralized to ∼pH 7.0 by the addition of 1 M Tris-HCl buffer.

### Enzyme-linked immunosorbent assay (ELISA)

Our earlier study (Ren et al., 2022) also describes the ELISA performed. The optical density was measured at wavelength 405 nm (OD_405_) by a microplate photometer (Multiskan™ FC, Thermo Fisher Scientific).

### Viruses

BIKEN Innovative Vaccine Research Alliance Laboratories (Osaka, Japan) provided the SARS-CoV-2 strain that contains the spike D614G mutation (DNA Data Bank of Japan [DDBJ]: accession no. LC644163), and in the present study we refer to this strain as strain D614G. Japan’s National Institute of Infectious Disease (Tokyo) provided the SARS-CoV-2 strains of the Pango lineage AY.122 (EPI_ISL_2158617) which we used as the Delta variant, BA.1.18 (EPI_ISL_7418017) which we used as the Omicron BA.1 variant, BA1.1 (EPI_ISL_7571618) which we used as the BA1.1 variant, BA.2 (EPI_ISL_9595859) which we used as the BA.2 variant, BA.2.75 (EPI_ISL_13969765), which we used as the BA.2.75 variant, and BA.5 (EPI_13241867), which we used as the BA.5 variant. We propagated the viruses by the infection of Vero E6 (TMPRSS2) cells in 2% FBS containing DMEM (Matsuyama et al., 2020) in order to create a stock of each virus (Furukawa et al., 2021).

### Plaque reduction neutralization test (PRNT)

To conduct a plaque reduction neutralization test (PRNT) and determine the neutralization percentage for each virus strain, we seeded Vero E6/TMPRSS2 cells (2□×□10^5^ cells/well) on 12-well plates (Corning) and cultivated them for 24 hr with 5% CO_2_ at 37°C. The cell monolayers were then washed one time with DMEM (without FBS). Each antibody diluted in DMEM (without FBS) was mixed with 100 plaque forming units (PFU) of SARS-CoV-2 and incubated for 1 hr at 37°C. We then added the virus–antibody mixture to the Vero E6/TMPRSS2 cells and cultured the cells for 1 hr with 5% CO_2_ at 37°C. After the removal of the inoculum, the infected cells were washed twice with PBS and incubated for 3–6 days at 37°C with 5% CO_2_ together with DMEM containing 2% FBS and 1.6% methylcellulose.

After the removal of the culture medium, PBS was used to wash the cells two times, and the cells were fixed with 80% methanol for 1 hr at room temperature. The remaining cells were stained with 1% crystal violet in 50% methanol for the visualization of plaques, which were counted manually. The ratio of neutralization was obtained by dividing the number of plaques obtained without antibody by the number of plaques obtained with antibody.

### Bio-layer interferometry

The affinity between each antibody and spike protein was measured by bio-layer interferometry (BLI) with a BLItz Biolayer Interferometer System (Sartorius, Goettingen, Germany).

### Ethics statement

The Kobe University Graduate School of Medicine’s Ethics Committee approved the collection and use of the COVID-19 patients’ blood samples (approval code: B200200). The patients’ written informed consent for this use was obtained.

## Notes

### Competing Interest Statement

The authors have declared no competing interest.

## References

Ai, J., Wang, X., He, X., Zhao, X., Zhang, Y., Jiang, Y., Li, M., Cui, Y., Chen, Y., Qiao, R., et al. (2022). Antibody evasion of SARS-CoV-2 Omicron BA.1, BA.1.1, BA.2, and BA.3 sub-lineages. Cell Host Microbe.

Cao, Y., Yisimayi, A., Jian, F., Song, W., Xiao, T., Wang, L., Du, S., Wang, J., Li, Q., Chen, X., et al. (2022). BA.2.12.1, BA.4 and BA.5 escape antibodies elicited by Omicron infection. Nature.

Dejnirattisai, W., Huo, J., Zhou, D., Zahradník, J., Supasa, P., Liu, C., Duyvesteyn, H.M.E., Ginn, H.M., Mentzer, A.J., Tuekprakhon, A., et al. (2022). SARS-CoV-2 Omicron-B.1.1.529 leads to widespread escape from neutralizing antibody responses. Cell 185, 467-484.e415.

Dong, J., Zost, S.J., Greaney, A.J., Starr, T.N., Dingens, A.S., Chen, E.C., Chen, R.E., Case, J.B., Sutton, R.E., Gilchuk, P., et al. (2021). Genetic and structural basis for SARS-CoV-2 variant neutralization by a two-antibody cocktail. Nat Microbiol 6, 1233–1244.

Furukawa, K., Tjan, L.H., Kurahashi, Y., Sutandhio, S., Nishimura, M., Arii, J., and Mori, Y. (2022). Assessment of Neutralizing Antibody Response Against SARS-CoV-2 Variants After 2 to 3 Doses of the BNT162b2 mRNA COVID-19 Vaccine. JAMA Netw Open 5, e2210780.

Furukawa, K., Tjan, L.H., Sutandhio, S., Kurahashi, Y., Iwata, S., Tohma, Y., Sano, S., Nakamura, S., Nishimura, M., Arii, J., et al. (2021). Cross-Neutralizing Activity Against SARS-CoV-2 Variants in COVID-19 Patients: Comparison of 4 Waves of the Pandemic in Japan. Open Forum Infect Dis 8, ofab430.

Hsieh, C.L., Goldsmith, J.A., Schaub, J.M., DiVenere, A.M., Kuo, H.C., Javanmardi, K., Le, K.C., Wrapp, D., Lee, A.G., Liu, Y., et al. (2020). Structure-based design of prefusion-stabilized SARS-CoV-2 spikes. Science 369, 1501–1505.

Kurahashi, Y., Furukawa, K., Sutandhio, S., Tjan, L.H., Iwata, S., Sano, S., Tohma, Y., Ohkita, H., Nakamura, S., Nishimura, M., et al. (2022a). Cross-neutralizing activity against Omicron could be obtained in SARS-CoV-2 convalescent patients who received two doses of mRNA vaccination. J Infect Dis.

Kurahashi, Y., Sutandhio, S., Furukawa, K., Tjan, L.H., Iwata, S., Sano, S., Tohma, Y., Ohkita, H., Nakamura, S., Nishimura, M., et al. (2022b). Cross-Neutralizing Breadth and Longevity Against SARS-CoV-2 Variants After Infections. Front Immunol 13, 773652.

Matsuyama, S., Nao, N., Shirato, K., Kawase, M., Saito, S., Takayama, I., Nagata, N., Sekizuka, T., Katoh, H., Kato, F., et al. (2020). Enhanced isolation of SARS-CoV-2 by TMPRSS2-expressing cells. Proc Natl Acad Sci U S A 117, 7001–7003.

Muik, A., Lui, B.G., Wallisch, A.K., Bacher, M., Muhl, J., Reinholz, J., Ozhelvaci, O., Beckmann, N., Guimil Garcia, R.C., Poran, A., et al. (2022). Neutralization of SARS-CoV-2 Omicron by BNT162b2 mRNA vaccine-elicited human sera. Science 375, 678–680.

Niwa, H., Yamamura, K., and Miyazaki, J. (1991). Efficient selection for high-expression transfectants with a novel eukaryotic vector. Gene 108, 193–199.

Ojima-Kato, T., Nagai, S., and Nakano, H. (2017). Ecobody technology: rapid monoclonal antibody screening method from single B cells using cell-free protein synthesis for antigen-binding fragment formation. Sci Rep 7, 13979.

Ren, Z., Nishimura, M., Tjan, L.H., Furukawa, K., Kurahashi, Y., Sutandhio, S., Aoki, K., Hasegawa, N., Arii, J., Uto, K., et al. (2022). Large-scale serosurveillance of COVID-19 in Japan: Acquisition of neutralizing antibodies for Delta but not for Omicron and requirement of booster vaccination to overcome the Omicron’s outbreak. PLoS One 17, e0266270.

Tan, C.W., Lim, B.L., Young, B.E., Yeoh, A.Y., Yung, C.F., Yap, W.C., Althaus, T., Chia, W.N., Zhu, F., Lye, D.C., et al. (2022). Comparative neutralisation profile of SARS-CoV-2 omicron subvariants BA.2.75 and BA.5. Lancet Microbe.

Tegally, H., Moir, M., Everatt, J., Giovanetti, M., Scheepers, C., Wilkinson, E., Subramoney, K., Moyo, S., Amoako, D.G., Baxter, C., et al. (2022). Continued Emergence and Evolution of Omicron in South Africa: New BA.4 and BA.5 lineages. medRxiv, 2022.2005.2001.22274406.

Tjan, L.H., Furukawa, K., Kurahashi, Y., Sutandhio, S., Nishimura, M., Arii, J., and Mori, Y. (2022). As well as Omicron BA.1, high neutralizing activity against Omicron BA.2 can be induced by COVID-19 mRNA booster vaccination. J Infect Dis.

Westendorf, K., Zentelis, S., Wang, L., Foster, D., Vaillancourt, P., Wiggin, M., Lovett, E., van der Lee, R., Hendle, J., Pustilnik, A., et al. (2022). LY-CoV1404 (bebtelovimab) potently neutralizes SARS-CoV-2 variants. Cell Rep 39, 110812.

Yamasoba, D., Kimura, I., Nasser, H., Morioka, Y., Nao, N., Ito, J., Uriu, K., Tsuda, M., Zahradnik, J., Shirakawa, K., et al. (2022a). Virological characteristics of the SARS-CoV-2 Omicron BA.2 spike. Cell 185, 2103–2115 e2119.

Yamasoba, D., Kosugi, Y., Kimura, I., Fujita, S., Uriu, K., Ito, J., and Sato, K. (2022b). Neutralisation sensitivity of SARS-CoV-2 omicron subvariants to therapeutic monoclonal antibodies. Lancet Infect Dis 22, 942–943.

